# Eye-movement related brain potentials during assisted navigation in real-world

**DOI:** 10.1101/2020.06.08.139469

**Authors:** Anna Wunderlich, Klaus Gramann

## Abstract

Conducting neuroscience research in the real world remains challenging because of movement- and environment-related artifacts as well as missing control over stimulus presentation. The present study demonstrated that it is possible to investigate the neuronal correlates underlying visuo-spatial information processing during real-world navigation. Using mobile EEG allowed for extraction of saccade- and blink-related potentials as well as gait-related EEG activity. In combination with source-based cleaning of non-brain activity and unfolding of overlapping event-related activity, brain activity of naturally behaving humans was revealed even in a complex and dynamic city environment.

## 1 INTRODUCTION

### Navigation Assistance Systems and Spatial Knowledge Acquisition

Navigating and orienting in our environment are fundamental aspects of every-day activities. Common navigation tasks vary with regards to the distance travelled and familiarity of the environment as they range from commuting to work, or grocery shopping to touristic trips, or long hikes to explore new areas. Increasingly, technology, i.e. navigation assistance systems, facilitate or even take over parts of these spatial orienting tasks. The frequent use of navigation aids, however, was shown to be associated with decreased processing of the environment and to be detrimental to the ability to successfully use spatial strategies when no navigation aid is available (Dahmani and Bohbot, 2020; Münzer et al., 2006; Ishikawa et al., 2008).

Furthermore, in previous studies we have shown that the use of commercial navigation instructions leads to a decrease of landmark knowledge, especially regarding knowledge of landmarks at decision points (Gramann et al., 2017; Wunderlich and Gramann, 2018; 2020). These studies further demonstrated the successful incidental acquisition of landmark and route knowledge when landmark-based rather than standard instructions were used. The experimental setups in these studies ranged from simulated driving through a virtual world Wunderlich and Gramann (2018) to interactive videos of walking through the real world or actual walking through the real-world (Wunderlich and Gramann, 2020). The results of Wunderlich and Gramann (2018) further showed changes in the event-related brain activity with cued recall of landmark pictures, which corresponded to the performance differences observed (Wunderlich and Gramann, 2018). Even though these studies provided new insights into spatial learning when assistance systems were used for navigation, they all addressed spatial knowledge acquisition *after* the assisted navigation phase providing no further insights into incidental spatial knowledge acquisition *during* navigation.

### Investigating Brain Activity During Navigation

To better understand spatial knowledge acquisition during assisted navigation, neuroscientific methods can be used. These methods allow for investigating human brain dynamics while navigators orient in their surroundings. However, established brain imaging methods require stationary laboratory setups and immobile participants to avoid movement-related artifacts from contaminating the signal of interest. Using new mobile brain imaging devices, in contrast, allows for recording human brain dynamics during active navigation and in the real world providing high ecological validity (Park et al., 2018).

Using light-weight EEG amplifiers with relatively high-density montages allows for investigating the neural dynamics underlying real-world navigation including natural interaction with a complex, dynamically changing environment and other social agents, as well as realistic visuals and soundscapes. However, mobile EEG recordings come with several problems. First, active movement through the real world is associated with increasing noise in the recordings (Gramann et al., 2014). The EEG collects data on the surface of the scalp that is the result of volume conducted brain and non-brain sources. The latter include biological sources (e.g., eye movement and muscle activity) as well as mechanical and electrical artifacts (e.g., loose electrodes, cable sway, electrical sources in the environment). A second problem lies in a multitude of external and internal events that are impossible to control but that are naturally present when the real world is used as experimental environment to investigate cognitive phenomena. Some of these events might provoke artifactual activity with respect to the phenomena of interest (e.g., startle responses to a car horn or suddenly appearing pedestrians). Finally, the real world might not provide the necessary quantity and timing of controlled external event presentations necessary for the analyses of event-related brain activity.

The first problem of inherently noisy data can be addressed by blind source separation methods such as independent component analyses (ICA, Bell and Sejnowski (1995), Makeig et al. (1996)). Removing non-brain sources from the decomposition allows for back-projecting only brain activity to the sensor level, using ICA as an extended artifact removal tool (Jung et al., 2000). The second problem, the multitude of random events, might be overcome by using an averaging approach of event-related potentials (ERPs) to average out EEG activity that was not related to the processes of interest. However, to do so, the third problem has to be solved and a sufficiently high number of meaningful events has to be found for event-related analyses (Luck et al., 2000).

### Extracting Events in Real-World Studies

The real world does not allow for a full control of the experimental environment leading to variations in visual or auditory stimuli perceived by participants. Even more problematic is the fact that tasks in the real world usually do not allow for a continuous repetition of the same or similar events of interest that can be responded to with a single button press. As a consequence, real-world experiments often offer only an insufficient number of controlled events for subsequent event-related analyses (e.g. the number of navigation instructions).

To overcome this problem, physiological non-brain activity captured in the mobile EEG can serve as a source for meaningful events for the analyses of ERPs as they are non-intrusive to the ongoing task (Bentivoglio et al., 1997). Thus, naturally occurring physiological events like eye blinks and saccades allow to parse the EEG signal into meaningful segments as they interrupt visual information intake (Stern et al., 1984jan; Berg and Davies, 1988; Kamienkowski et al., 2012). Saccades suppress visual information intake starting 50 ms preceding saccade-onset as well as during the saccade. Thus, each fixation following a saccade represents onset of visual inormation intake. Event-related potentials using saccades can be related to either saccade onset, peak velocity, or saccade offset, with the latter being equivalent to the fixation-related potential (fERP). Saccade-related brain potentials (sERP) were used in many previous studies (Gaarder et al., 1964dec;Rämä and Baccino, 2010), especially in research investigating reading and text processing (Marton and Szirtes, 1988; Baccino, 2012; Dimigen et al., 2011) or visual search (Ossandón et al., 2010; Kaunitz et al., 2014; Kamienkowski et al., 2018).

Blinks produce a longer interruption of the visual input stream (for a review see Stern et al. (1984jan)) with three different functions underlying a potential blink generation. First, in order to keep efficiency of the visual input channel high, and reduce the number and time of interruptions of the visual information stream, they are combined with other eye-movements (Evinger et al., 1994). Second, blinks likely occur after a period of blink suppression, e.g. during attention allocation, or when the processing mode changes. Thus they can mark the end of processing steps (Stern et al., 1984jan). Third, in very structured tasks using for example stimulus-response pairs blinks show a temporal relationship to those events (Stern et al., 1984jan).

Like saccade- and fixation-related eye movements, blinks have been shown to allow for extracting event-related potentials (bERP). The bERP was shown to be sensitive to parameters of the experimental environment or of the current task (Berg and Davies, 1988; Wascher et al., 2014). The longer preceding pause of incoming information might make bERPs more similar to ERPs of complex scene-stimuli than fERPs. In addition, the likely timing of blinks at the end of processing steps qualifies the bERPs during natural viewing a valuable source of insight about visual information processing and underlying cognitive processes.

### Analyzing Mobile EEG Recordings in the Real World

In this paper, we describe a means to deal with the issues arising from collecting mobile EEG during an ongoing task in the real world. We show how both blink- and saccade-related potentials, alongside with gait-related activity evoked during mobile EEG recordings in uncontrolled real-world environments can be extracted from IC activation and how these events can subsequently be analyzed to extract meaningful information from the ongoing brain activity.

In the present study, this approach was used to investigate the human brain activity underlying visual processing of the environment during navigation that was supported by landmark-based navigation instructions. We recorded and analysed brain activity in the real world while participants navigated through the city of Berlin receiving either standard or landmark-based navigation instructions. We expected differences in visual processing of the environment dependent on the navigation instruction condition reflecting a general increase in the awareness for surrounding information relevant to navigation in the environment as shown in the brain activity of the cued-recall task in Wunderlich and Gramann (2018). This was investigated using blink and saccade-related potentials extracted during different periods of the navigation task (during navigation instructions versus times where participants were simply walking straight segments of the route). A general increased awareness would reveal differences between navigation instruction conditions in both navigation phases. Contrasting, an occasional increased awareness to environmental features during the provision of navigation instructions would only reveal differences between the groups in the navigation instruction phase.

## 2 MATERIALS AND METHODS

### 2.1 Participants

The data of 22 participants (eleven women) was analyzed with eleven participants in each navigation instruction condition. Their age ranged from 20 to 39 years (*M* = 27.4, *SD* = 4.63 years). Participants were recruited through an existing database or personal contact and received either 10 Euro per hour or course credit. All had normal or corrected to normal vision and gave informed consent prior to the study which was approved by the local research ethics committee of the Institute for Psychology and Ergonomics at the Technische Universität (TU) Berlin. Before the main experiment, participants filled out an online questionnaire to determine if they were familiar with the area where the navigation task would take place as used in Wunderlich and Gramann (2020). After navigating the route, participants were again asked whether they had been familiar with the navigated route. In case of familiarity, they were instructed to rate the known percentage of the route. In case participants stated to have been familiar with more than 50% of the route, they were excluded from the second part of the experiment and data analysis. In the final sample of 22 participants, familiarity ratings ranged from 0 to 40% (*M* = 9.52%, *SD* = 12.2%).

### 2.2 Study Design and Procedure

The experiment consisted of two parts and lasted approximately three hours in total. In the first part, participants walked a predefined route through the district of Charlottenburg, Berlin in Germany, using an auditory navigation assistance system. In the second part, directly after the navigation task, participants were transported back to the Berlin Mobile Brain/Body Imaging Lab (BeMoBIL) at TU Berlin to run different spatial tests. Participants had not been aware that they would be tested on the environment after the navigation task.

During the navigation task, participants followed the auditory navigation instructions to navigate along a predefined, unfamiliar route with two groups receiving either standard or landmark-based navigation instructions. Based on previous results (Gramann et al., 2017; Wunderlich and Gramann, 2018; 2020), landmark-based instructions referenced a landmark at each intersection and provided more detailed information about this landmark. This condition will be referred to as *long* instruction condition in the following. One example navigation instruction for this long condition was ‘Turn left at the UdK. The UdK is the biggest University of Arts in Europe.’ This contrasted with the *standard* navigation instruction condition that used instructions like the ‘Turn left at the next intersection.’ In the instructions for the navigation task, it was pointed out to the participants that they should follow the auditory turn-by-turn instructions and be aware of other traffic participants, especially while crossing streets. Furthermore, in case they feel lost, they were asked to stop and turn to the experimenter who was always shadowing the participant. The presence of the experimenter ensured the participant’s safety as well as the correct course of the route as the experimenter manually triggered the auditory navigation instructions using a browser-based application on a mobile phone. The participant received the auditory navigation instructions by Bluetooth in-Ear headphones at predefined trigger points in the environment. After walking for approximately 40 min, the participants arrived at the end of the route. Their they were instructed to rate their subjective task load during navigation using the National Aeronautics and Space Administration Raw Task Load Index (NASA-RTLX, Hart (2006), Hart and Staveland (1988)). Additionally, they filled in three short questions regarding their prior knowledge of the route.

The second part of the experiment took place at the BeMoBIL. There, the first task was to draw a map of the route and secondly solve a cued-recall task. In the latter task, landmark pictures were given as cues and the required response included the respective route direction. The randomly shown landmarks had been either located at intersections (and mentioned in the landmark-based navigation instructions) or at straight segments of the route, or they were similar in appearance but not part of the previously navigated route. At the end, demographic data as well as individual navigation habits, and subjective spatial ability ratings were collected.

### 2.3 Electroencephalography

#### EEG Data Collection

The EEG was recorded continuously during the navigation task and the subsequent laboratory tests using an elastic cap with 65 electrodes (eego, ANT Neuro, Enschede, Netherlands). Electrodes were placed according to the extended 10% system (Oostenveld and Praamstra, 2001). All electrodes were referenced to CPz and the data was collected with a sampling rate of 500 Hz. One electrode below the left eye was used to record vertical eye movements. Time synchronization and disk recording of the EEG data stream and the event marker stream from the mobile application and task paradigm was done using Lab Streaming Layer (LSL, Kothe (2014)).

#### EEG Data Processing

For EEG data processing the Matlab toolbox EEGLAB was used (Delorme and Makeig, 2004). The raw EEG data of both the navigation phase and the cued-recall task were high pass filtered at 1 Hz, low pass filtered at 100 Hz using the EEGLAB filter function *eegfilter(),* and subsequently resampled to 250 Hz (see Figure 1, left column). The pre- and post-task phases of the EEG data were removed. Afterwards, the two separate datasets of each participant were merged into one dataset and channels that were subjectively judged as very noisy, flat, or drifting were manually rejected (M = 3.79, *SD* = 1.77, *Min* = 1, *Max* = 8). Continuous data cleaning was applied twice using the *pop_rejcont()* function for frequency limits from 1 to 100 Hz and default settings for all other parameters^1^. Once before and once after, the rejected channels were interpolated using a spherical spline function and the data was re-referenced to the average reference. The cleaning before interpolation and average reference was targeting at removal of artifacts on a single channel level. This way, the inflation of their impact to more channels was prevented when re-referencing to average reference. The second continuous cleaning was applied to remove remaining artifactual data.

**FIGURE 1.**
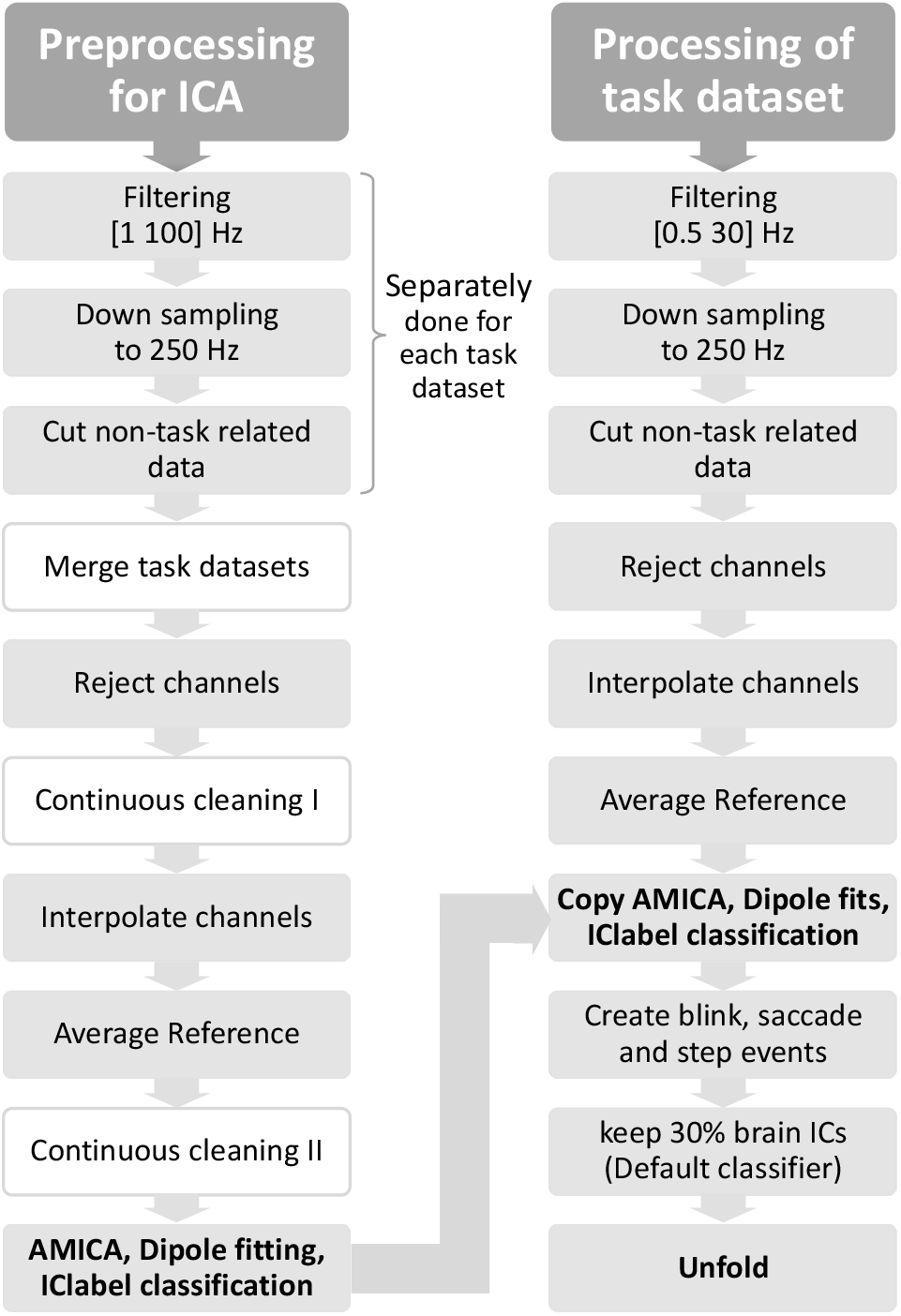
EEG data processing from raw data to ICA computation (left) and from raw task data to the use of the unfold toolbox (right). Additional analysis steps in the ICA preprocessing have white boxes to emphasize the otherwise parallel processing.

Subsequently, the data was submitted to independent component analysis (ICA, Makeig et al. (1996)) using the Adaptive Mixture ICA (AMICA, Palmer et al. (2011)). The resultant independent components (ICs) were localized to the source space using an equivalent dipole model as implemented in the dipfit routines (Oostenveld and Oostendorp, 2002). Finally, the resultant ICs were classified as being brain, muscle or other processes using the default classifier of IClabel (Pion-Tonachini et al., 2019).

The original sensor data was preprocessed using identical processing steps as described above save different filter frequencies and no time domain data cleaning (see Figure 1, right column). The respective weights and sphere matrices from the AMICA solution were applied to the preprocessed navigation dataset. In addition, the equivalent dipole models and IClabel classifications for each participant and IC were transferred to the respective task dataset allowing for the extraction of events based on the complete duration of the task.

#### Event Extraction from IC Activation

The event extraction is summarized in Figure 2. Blinks were identified using one IC from the individual decompositions that reflected vertical eye movements as described in Lins et al. (1993). In case of more than one candidates for the vertical eye IC, the one showing better signal to noise ratio for blink deflections and/or less horizontal eye movement was chosen based on subjective inspection. For detecting blinks, the associated component activation time course was filtered using a moving median approach (window size of twenty sample points equaling 80 ms). Moving median approaches smooth without changing the steepness of the slopes in the data (Bulling et al., 2010). To allow for automated blink peak detection, all individual ICs time courses were standardized to a positive peak polarity. Peak detection was done using the MATLAB function *findpeaks()* applied to the filtered IC activation. Parameters used were a *minimal peak distance* of 25 sample points (100 ms) to avoid directly following blinks to be selected. Further, peaks were restricted to a *minimal peak width* of 5 (20 ms) and *maximal peak width* of 80 sample points (320 ms) to suppress potential high-amplitude artefacts or slow oscillations from being counted as a blink. The following two parameters were automatically defined for each dataset individually to take care of interindividual differences in the shape of the electrical signal representing a blink: the 90-percentile of the filtered activation data was applied to define a threshold of *minimal peak prominence*. This parameter ensured the successful separation of detected peaks from the background IC activity. For the absolute *minimal peak height,* a threshold was defined using the 85-percentile of the filtered activation data. For each peak location, an event marker with the name ‘blink’ was created in the EEG dataset at the time point of maximum blink deflection.

**FIGURE 2.**
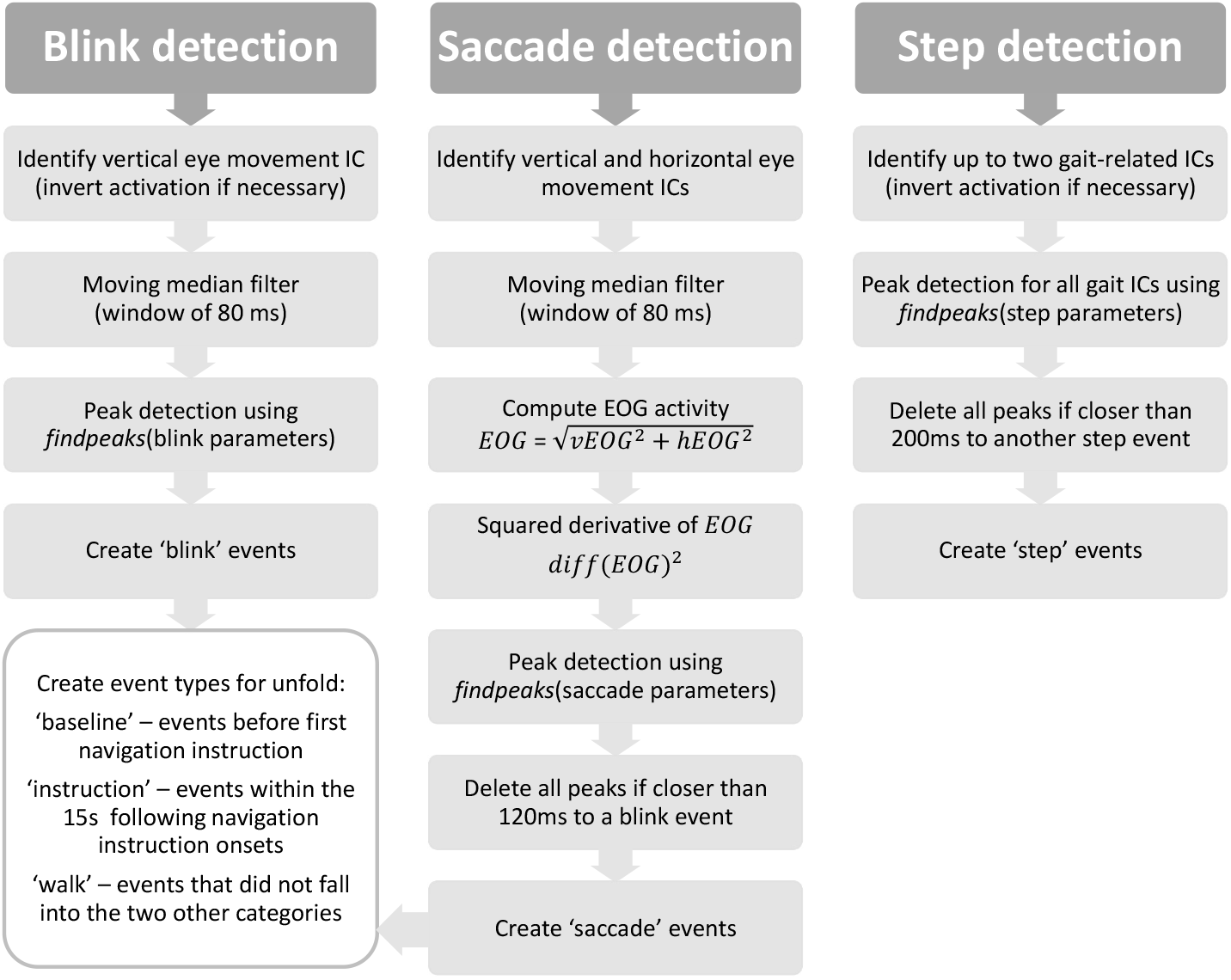
Analysis steps for the extraction of events from the respective IC activation(s): blink (left), saccade (middle), and step events (right). The respective parameters for the findpeaks() functions can be found in the text.

Saccades were identified using two ICs from the individual decompositions that reflected vertical and horizontal eye movements, respectively according to Lins et al. (1993). Vertical eye-movement ICs were the same as used for blink detection. For the horizontal eye-movement ICs, the IC with the characteristic scalp map and rectangular activity of horizontal eye movements was chosen based on subjective inspection. The associated component activation time courses were filtered using a moving median approach (window size of 20 sample points equaling 80 ms). The electrooculogram (EOG) activity was calculated using the root mean square of the smoothed time courses (Jia and Tyler, 2019). For saccade maximum velocity detection, the first derivative was taken and squared to increase the signal-to-noise-ratio. The function *findpeaks()* was applied on the squared derivative of the EOG activation. Parameters used were a *minimal peak distance* of 25 sample points (100 ms). Peaks were restricted to a *minimal peak width* of 1 (4 ms) and *maximal peak width* of 10 sample points (40 ms). The 90-percentile of the squared derivative of the EOG was applied for *minimal peak prominence* as well as for the *minimal peak height* threshold. In case identified peaks were closer than 30 sample points (120 ms) to a blink event, these peaks were not taken for saccade event extraction to avoid taking saccades into account that appeared during eyes closed periods. For each of the remaining peaks, an event marker called ‘saccade’ was created in the EEG dataset at the time point of maximum saccade velocity (middle of the saccade).

Gait-related activity was identified using up to two ICs from the individual decompositions that reflected steps in the gait cycle as described previously (Knaepen et al., 2015; Kline et al., 2015; Maidan et al., 2019). No filtering or smoothing was applied to the associated component activation time courses showing a pronounced waveform at approximately 2 Hz. Inverting of time courses for some ICs was done to align peak amplitudes on top of the slow wave maxima. To extract single steps of the gait cycle, *findpeaks()* was applied to both IC activations consecutively. Peaks were restricted to a *minimal peak width* of 5 (20 ms) to take advantage of the high frequency part and *maximal peak width* of 150 sample points (600 ms) to detect the slow wave peaks. *Minimal peak distance* was set to 100 sample points (400 ms) to avoid that the high frequency part and the slow wave peak were both used for event extraction. The 80-percentile of the IC activation time course was applied for *minimal peak prominence* and as threshold for *minimal peak height.* In case of step events identified in the two ICs being closer than 50 sample points (200 ms) one of the respective events were not taken in account for event generation. For each remaining detected peak, an event marker named ‘step’ was created in the EEG dataset at the time point of putting full weight on the respective foot.

Afterwards every dataset was visually checked to validate blink, saccade, and gait events. For this, it was checked whether the events marked in EEG activity were representative for the respective body movement as reported before in the literature (Lins et al., 1993; Kline et al., 2015). To enable the comparison of blink- and saccade-related brain activity in different phases of the navigation task, we included a subordinate event type. Either ‘baseline’ in case the event appeared before the first navigation instruction and was thus unaffected by the navigation instruction conditions, or ‘instruction’, in case the event took place within the 15 seconds following the onset of a navigation instruction. The category ‘walk’ was used as event type for all other remaining events in the navigation phase. The number of events in each category can be seen in Table 1.

**TABLE 1.** Number of blink, saccade, and step events for all participants and separated by navigation instruction condition and navigation phase.

|  |  | Number blink events |  |  | Number saccade events |  |  | Step |
| --- | --- | --- | --- | --- | --- | --- | --- | --- |
|  |  | Baseline | Instruction | Walk | Baseline | Instruction | Walk | All |
| **Overall** | *M* | 198 | 203 | 1310 | 606 | 730 | 4347 | 4105 |
|  | *SD* | 136 | 79 | 612 | 258 | 163 | 1144 | 952 |
|  | *Min* | 68 | 89 | 384 | 184 | 366 | 1893 | 1762 |
|  | *Max* | 572 | 368 | 2907 | 1307 | 1023 | 6808 | 6878 |
| **Standard** | *M* | 214 | 213 | 1385 | 652 | 666 | 4386 | 4216 |
| **instruction** | *SD* | 142 | 81 | 592 | 224 | 113 | 1104 | 1033 |
| **condition** | *Min* | 80 | 99 | 693 | 294 | 430 | 3184 | 3131 |
|  | *Max* | 572 | 355 | 2730 | 996 | 857 | 6808 | 6878 |
| **Long** | *M* | 183 | 193 | 1236 | 559 | 794 | 4307 | 3994 |
| **instruction** | *SD* | 128 | 76 | 623 | 281 | 180 | 1181 | 850 |
| **condition** | *Min* | 68 | 89 | 384 | 184 | 366 | 1893 | 1762 |
|  | *Max* | 525 | 368 | 2907 | 1307 | 1023 | 5818 | 5166 |

M, mean; SD, standard deviation; Min, Minimum; Max, Maximum.

#### Source-based EEG Data Cleaning

Subsequently, all ICs with a classification probability lower than 30 % in the category *brain* were removed from the dataset and the data was back-projected to the sensor level. This way the number of ICs per participant was reduced to *M* = 13.3 ICs (SD = 4.50 ICs, *Min* = 5 ICs, *Max* = 22 ICs). Considering the instruction conditions, this IC reduction did not lead to unbalanced numbers of ICs between the two instruction condition groups (standard navigation instructions: *M* = 13.1 ICs, *SD* = 5.12 ICs, *Min* = 5 ICs, *Max* = 22 ICs; long navigation instructions: *M* = 13.5 ICs, *SD* = 4.74 ICs, *Min* = 6 ICs, *Max* = 18 ICs).

#### Unfolding of Event-Related Activity

The last data processing step on the single subject level was the application of the unfold toolbox to the continuous data (Ehinger and Dimigen, 2019). This toolbox allows for a regression-based separation of overlapping event-related brain activity. As the extracted eye and body movement events in the navigation task overlapped with each other (Dimigen et al., 2011) and/or were temporally synchronized for some participants, it is a valuable tool to consider and control for overlapping ERPs and those individual differences.

Following the published analysis pipeline of Ehinger and Dimigen (2019), we defined a design matrix with blink, saccade and step events and 64 channels. We included the categorial factor *navigation phase* (baseline, instruction, walk) for the blink and saccade events into the regression *formula: y ~1 + cat(navigation phase).* For the step events, we only computed the intercept: *y ~1.* After applying the continuous artifact detection of the unfold pipeline and exclusion set to *amplitude threshold* of 80 μV, we time-expanded the design matrix according to the *timelimits* of −500 ms and 1000 ms referring to the event timestamp. Afterwards, we fitted the general linear model and extracted the intercept and beta values considering similar to Wascher et al. (2014) −500 ms to −200 ms for baseline correction.

While the blink and saccade related potentials were considered as informative for the analysis of visual information processing during navigation, the step events were only used to control for their individual impact on the blink and saccade related potentials. The intercept and beta values of the general linear model built the basis for a comparison between participants. Unfolded event-related potentials for all event types and their differences to the respective baseline events were plotted for a subset of 34 electrodes. The ERPs of participants within one navigation instruction condition and navigation phase were averaged and plotted alongside with twice the standard error of the mean (SEM).

### 2.4 Statistical Analysis

Statistical analysis was performed for the interaction of navigation instruction condition and navigation phase for the blink- and saccade-related potentials. Difference plots of the ERPs for both navigation phases (*instruction* and *walk* minus *baseline* were investigated to find time windows revealing significant differences between the navigation instruction conditions after the single trial baseline ending at −200 ms. To define statistical significance between the unpaired values, the EEGLAB function *statcondfieldtrip()* was used. Due to this, a 10000 fold permutation testing was applied followed by a cluster-based correction for family-wise error. If the returned probability corrected two-tailed p-Value was below .05 the sample was marked as statistically significant.

## 3 RESULTS

Blink- and saccade-related potentials were extracted from the EEG data during assisted navigation through the real world. The aim was to investigate the impact of landmark-related navigation instructions on ongoing brain activity underlying visual information processing and learning of environmental features. The present study aimed at establishing a principled pipeline of data processing and event extraction that allows for analyses of human brain dynamics in the real world addressing movement-related artifacts as well as EEG activity induced by uncontrollable environmental factors.

Essential processing steps were the spatial filter computation using ICA, removal of ICs reflecting non-brain activity, and unfolding of overlapping events. The resulting unfolded event-related potentials can be seen in Figure 3 for saccades and in Figure 4 for blink events. ERPs were plotted for a subset of 34 electrodes distributed over the whole scalp using baseline correction from −500 to −200 ms. Separate lines represent the different navigation phases (line style) and navigation instruction conditions (grey intensity).

**FIGURE 3.**
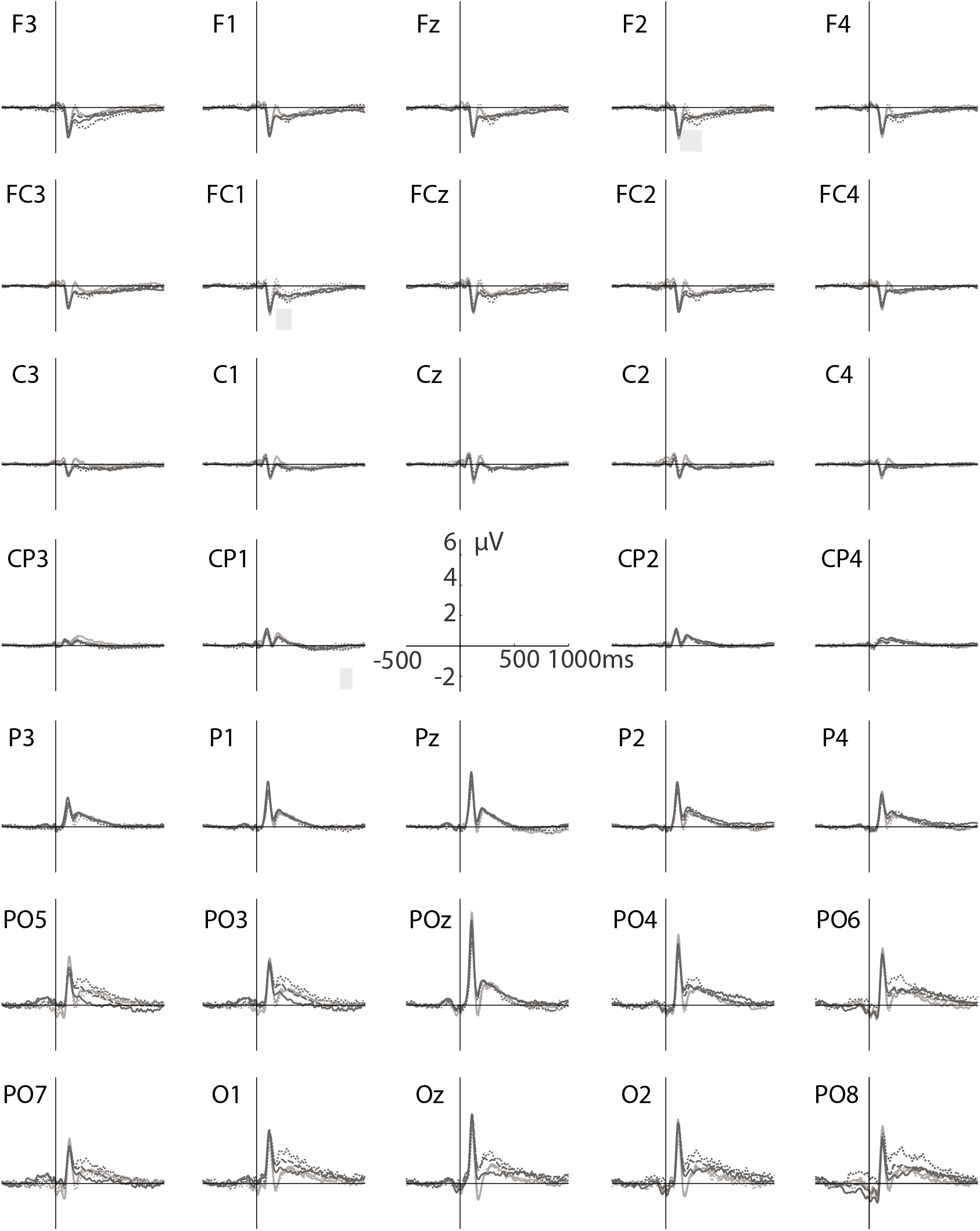
Saccade-related potentials displayed over the whole scalp averaged for each navigation instruction conditions separately (light grey - standard, dark grey - long), instruction phase (solid line) versus walk phase (dashed line), and baseline phase (dotted line). The grey rectangular areas represent time windows where significant differences between the navigation instruction conditions were found in the beta values for walk phase (light grey). There were no time windows with significant differences in the instruction phase. Positivity is plotted upwards.

**FIGURE 4.**
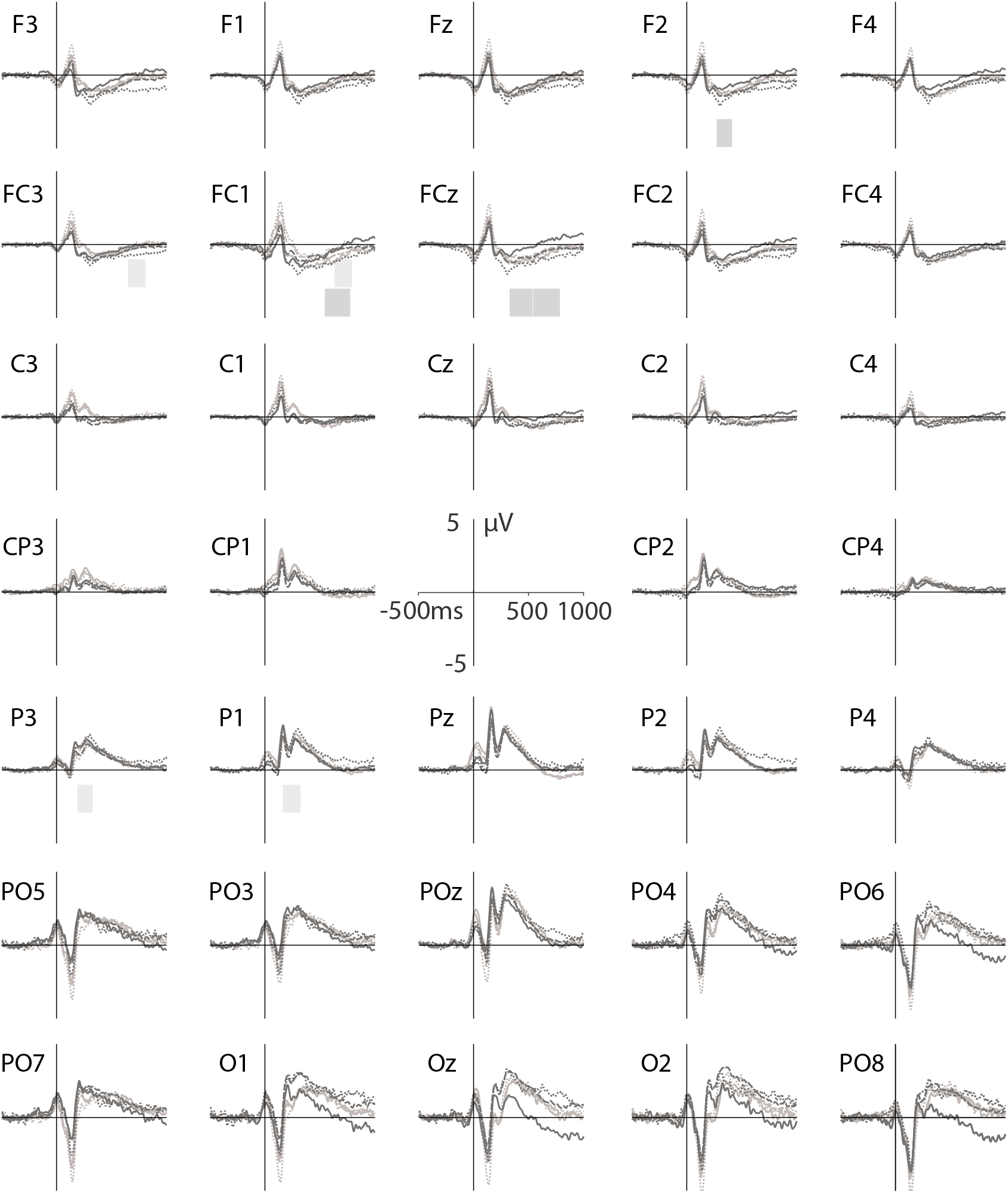
Blink-related potentials displayed over the whole scalp averaged separately for each navigation instruction condition (light grey - standard, dark grey - long), instruction phase (solid line), walk phase (dashed line), and baseline phase (dotted line). The grey rectangular areas represent time windows where significant differences between the navigation instruction conditions were found in the beta values for the instruction phase (dark grey) and walk phase (light grey). Positivity is plotted upwards.

### 3.1 I Saccade-related potentials

The order and polarity of sERP components was comparable across navigation instruction conditions and navigation phases. Amplitudes increased gradually from frontal to occipital leads as well as from lateral leads towards the central midline. When describing the revealed potential shape, we label the peaks using the polarity and a rounded to a multiple of 50 ms as well as names established in sERP and fERP literature.

The spline potential or P0 was revealed at the majority of electrodes about 10 to 20 ms preceding to the extracted saccade event.

At frontal leads, a first negative component (N100) following the saccade event became visible at 80 ms peaking at 120 ms with an amplitude maximum of-2 μV at Fz. After the N100, apositive component was observed peaking at 160 ms after the saccade event. Subsequently the potential slowly returned to the baseline without further identifiable components. The peak amplitude of the N100 decreased from frontal leads towards more central electrodes while the N100-peak on- and offset form small elevations.

At parietal and occipital leads a pronounced P100 or lambda response was elicited peaking at 110 ms. The P100 had the highest amplitude at POz with 6 μV. Following the P100, a second negative component (*N150)* with a minimum at 160 ms became visible. After this N150, a second positive component (*P200)* evolved peaking around 200 ms at Pz and POz and around 300 ms at Oz. Afterwards, the potential slowly returned to its baseline without further components. The parietal P100 was slightly more pronounced over the right hemisphere when compared to left hemispheric leads.

### 3.2 Blink-related potentials

The bERP revealed a different pattern of peaks compared to the sERP that showed instruction dependent amplitude differences. The order and polarity of components was the same across ERPs for different navigation instruction conditions and navigation phases. All components were more pronounced at the central line compared to right or left hemispheric peripheral electrodes.

From frontal to central sites, the bERP revealed a first negative component when the eyes were completely closed (*N0*). After this first N0, the frontal bERP activity raised to the first positive component (*P150)* reaching a maximum of about 2 μV around 140 ms after the blink at frontal leads with decreasing amplitudes towards lateralized leads. A second negative component was observed around 200 ms (*N200)* which was most pronounced at frontal leads with −1 μV and about 0 μV at central leads. A second but relative small positivity peaked around 250 ms (*P250*), followed by a second negative component around 300 ms (N300). This N300 was not as much pronounced as the previous N200 and remained at this level for about 200 ms or slowly arose towards baseline level. After a small discontinuity at about 500 ms the activity slowly ascended further towards 0 μV. This late negative potential around 600 ms will be referred to as the late negative level (*LNL*).

Parietal to occipital leads revealed a small elevation peaking at time point zero (*P0*) as a counterpart to the frontal N0. This was followed by a negativity (*N100*) with increasing amplitudes and latencies from parietal to occipital sites (Pz: 0.8 μV, 110 ms; POz: −0.5 μV, 120 ms; Oz: −2.5 μV, 130 ms). Event-related potentials show the sum activity of possibly temporally overlapping components with same or opposite polarity. Assuming a likewise attenuation of the positive component at Oz by the previous strongly pronounced negative component, the time shift between electrode sites of the latter component was also revealed in the following positive Peak (*P150*) as well as in the remaining amplitude. The maximal amplitude was elicited at Pz with 4 μV and a latency of 160 ms. At Oz the latency of the remaining peak of the positivity was located around 200 ms post event (*P200*) and reached 0 μV in the standard as compared to 2 μV in the long navigation instruction condition. This instruction dependent variation was visible in all navigation phases. After the P150, a second negative component (*N250)* peaked at about 1.5 μV with a latency of 230 ms at electrodes Pz and POz. At Oz, the negativity was temporally aligned with the one at Pz and POz and the peak-to-peak difference to the previous positivity constant for both experimental groups. Another positive component followed with a peak around 300 ms (*P300)* most pronounced at POz with about 3 μV. Reaching P300 the difference between the navigation instruction conditions at Oz levelled out. Finally, also at parietal to occipital leads a small discontinuity at approximately 600 ms was noticeable combined with an increased flattening of the slope compared to before.

### 3.3 Differences Between Navigation Instruction Conditions

The results of the statistical analysis are displayed as grey bars in Figures 4 and 3. The ERPs were tested samplewise for significant differences between the navigation instruction conditions and separately for the walk and the instruction phase. In order to do so, the event-related activity of each participant was controlled for the respective event-related activity before the first navigation instruction (baseline phase) by testing the beta values provided by the unfold toolbox (see Figure 5 top row).

**FIGURE 5.**
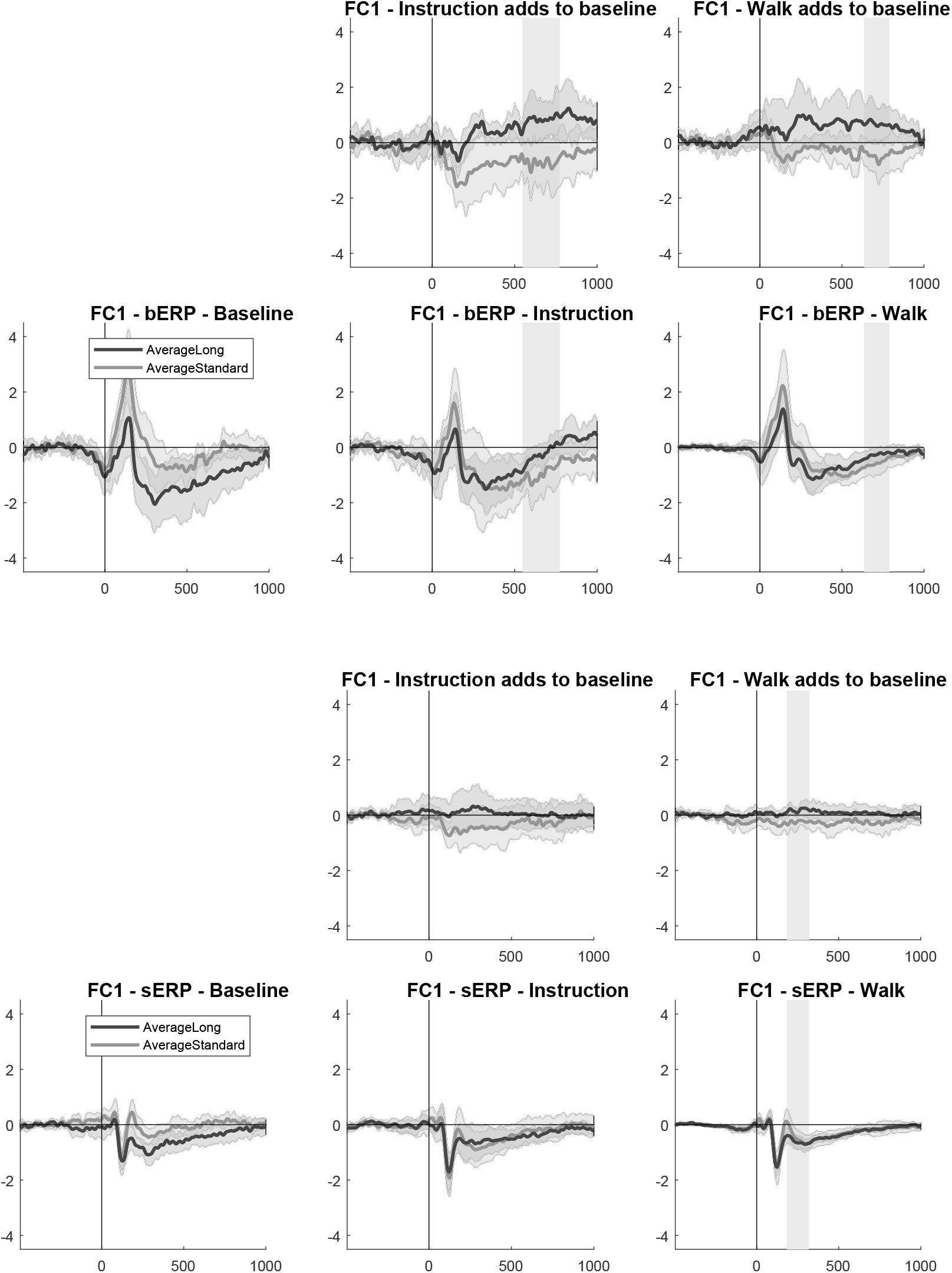
Blink-related potentials (Upper panel) and saccade-related potentials (Lower panel) for all three navigation phases. Above the difference plot of either the navigation or the walk phase and the baseline is plotted. The grey background areas represent time windows where significant differences between the navigation instruction conditions were found in the beta values. Positivity is plotted upwards.

Time windows containing significant samples surviving the family-based error correction were plotted as grey bars underneath the respective ERPs (using light grey for differences in walk and dark grey for instruction phase differences compared to baseline phase in Figures 4 and 3 and as simple grey background in Figure 5).

Blink-related potentials differed dependent on the navigation instruction conditions, irrespective of baseline differences, at F2, FC1, and FCz for the navigation instruction phase and at FC3, FC1, P3, and P1 for the walk phase. In the saccade-related potentials, significant differences were observed only for the walk phase at F2, FC1, and CP1.

The timing of the significant differences and their accompanying peaks in the event-related potentials are listed in Table 2. Additionally, the respective navigation instruction condition that showed higher absolute event-related activity in this time window is listed in the last column. Saccade-related activity differed during the baseline phase of the task dependent on the two groups of navigation instruction conditions while no such differences were observed during the instruction or the walk phase.

**TABLE 2.** Significant time windows in the blink- and saccade-related potentials.

| Event | Phase | Electrode |  |  | Time in ms<br>start-end | Peak in ERP | Instruction condition<br>with higher values |
| --- | --- | --- | --- | --- | --- | --- | --- |
|  |  | Left | Center | Right |  |  |  |
| Blink | Instruction |  |  | F2 | 280 - 408 | N300 | long |
|  |  | FC1 |  |  | 552 - 772 | LNL | long |
|  |  |  | FCz |  | 336 - 532 | N300 | long |
|  |  |  | FCz |  | 548 - 780 | LNL | long |
|  | Walk | FC3 |  |  | 656 - 804 | LNL | qualified by BL diff |
|  |  | FC1 |  |  | 640 - 788 | LNL | long |
|  |  | P3 |  |  | 196 - 324 | P300 | qualified by BL diff |
|  |  | P1 |  |  | 172 - 320 | N250, P300 | standard |
| Saccade | Walk |  |  | F2 | 140 - 328 | N300 | qualified by BL diff |
|  |  | FC1 |  |  | 188 - 320 | N300 | qualified by BL diff |
|  |  | CP1 |  |  | 776 - 880 | LNL | qualified by BL diff |

BL diff, difference between the instruction conditions in the baseline phase; ERP, event-related potential; LNL, late negative level.

## 4. DISCUSSION

This study investigated the neural basis of visuo-spatial information processing of pedestrians navigating through a real city guided by an auditory navigation assistance system. The auditory navigation instructions were either standard navigation instructions providing turning instructions referring to the next intersection or landmark-based navigation instructions referring to a salient object at the upcoming intersection and providing explanatory information about the landmark object. The changes in visual information processing during assisted navigation in the real-world were investigated by extracting saccade- and blink-related potentials from the recorded mobile EEG data while controlling for gait-related activity. This way a sufficient number of events for analyzing event-related brain activity in an otherwise uncontrolled real world environment was possible. The resulting blink-based ERPs proved to be sensitive to the experimental manipulations of navigation instructions.

### 4.1 Saccade-related potentials

The analyses of saccade-related potentials revealed clear peaks that had been reported previously, including the spline potential, the lambda wave and a clear posterior P2 component.

#### Early Evoked Components

The spline potential represents the saccade onset and was revealed across the scalp 10 to 20 ms before the extracted saccade event representing peak velocity of the saccade.

The P100 was most pronounced at POz. This is in line with the lambda response recorded at posterior leads in sERPs and fERPs. The lambda response is the best investigated saccade-related component and sensitive to properties of the visual stimulus like luminance or contrast (Gaarder et al., 1964dec; Kazai and Yagi, 2003; Dimigen et al., 2011; Kaunitz et al., 2014). This sensitivity of the lambda response to properties of incoming visual information is comparable to the P1 in stimulus-evoked ERPs. Furthermore, the underlying dipole producing the lambda response was shown to be located very close to the dipole forming the P1 (Kazai and Yagi, 2003). Thus, P1 and lambda response seem to be elicited by the same perceptual process (Kaunitz et al., 2014). Due to the dipolar activity pattern of this component, it is likely that the anterior N100 is the negative counterpart of the same activity source conveyed by volume conduction (Kamienkowski et al., 2012).

#### Later Evoked Components

The subsequent P2 at posterior leads was shown to be sensitive to processing of context information (Marton and Szirtes, 1988) and semantic meaning of text information (Simola et al., 2009). Simola et al. (2009) showed a right hemispheric dominance of the P2 component when processing semantically meaningful information versus non-words. In visual search, the P2 at Pz was related to a decreased amplitude for fixations of targets compared to distractors (Kamienkowski et al., 2012). In a later time window starting at 380 ms Kamienkowski and colleagues showed a positive component for targets only at frontal leads.

The statistical differences between the navigation instruction conditions shown in the sERPs of the walk phase at frontal leads are earlier than the frontal positive component shown in relation to targets in visual search (Kamienkowski et al., 2012). Additional, it seems likely that a difference in the baseline sERP levelled out with increasing time on task. There were no differences between the navigation instruction conditions during the instruction phase. Thus, this data revealed no saccade-related evidence for a visual search triggered by landmark references.

Achieving high ecological validity by recording EEG in mobile participants in the real world increased movement-related artifacts and variability in behavioral and cognitive strategies used to solve the task which. As a consequence, the high ecological validity renders the interpretation of the results more difficult (Parada, 2018).

### 4.2 Blink-related potentials

More pronounced frontal P100 and parietal P300 components in the bERP compared to the sERP point to a difference in information processing triggered by blinks. Blink-related activity revealed a stronger frontal as well as parietal activity with the latter most likely reflecting aspects of visuo-spatial information processing. Compared to previous studies, the order and polarity of the observed components 3.2 were roughly comparable to the results reported by Berg and Davies (1988) and Wascher et al. (2014) even though there were differences especially regarding negative components of the potential. The smoother appearance of the presented bERPs compared to Wascher et al. (2014) is likely due to the additional EEG data cleaning and processing steps in the present study.

A direct comparison of the peaks observed in the present study and those reported by Wascher et al. (2014) hints at regional differences between bERP components dependent on the task at hand. As a consequence, we labelled the bERP components in the present study according to their polarity and rounded latency after the blink event.

#### P0/N0

Opposite polarities of the early ERP peaks at anterior versus posterior electrode sites with most pronounced amplitudes over the occipital lobe at the time point of the blink likely reflect volume conducted activity of a radial source in or near the visual cortex. These early peaks evolve parallel with the blink and demonstrate inverse polarity compared to the maximum blink amplitude that served for event-extraction. Since the EOG activity was removed from the EEG data by removing the related independent components, this component is unlikely related to eye or lid movement. Rather, activation of the visual cortex associated with blinks could be a candidate underlying this evoked activity. A functional Magnet-Resonance-Tomography study, Tsubota et al. (1999jul) showed brain activity in the anterior areas of the occipital cortex when blinking in full darkness, which seems to be related to the control of the blink-movement.

#### N100/P150

Latency differences in the N100 at posterior leads andtheP150 at anterior leads speak against the same cortical source underlying these two components. The maximal amplitude and peak width of the N100 at Oz speaks for a component originating in the occipital cortex reflecting sensory processing of incoming visual information. In a very simple experiment of Berg and Davies (1988) light sensitivity was already shown in the early peaks of the bERPs at central and posterior electrodes pointing towards a very basic role of these components in visual information processing.

In general, a comparison of the bERPs components with established visual evoked ERP components like the posterior P1/N1 complex shows several deviations in morphology and latency. A reason for the observed latency difference could be that the current study used the blink-related amplitude maximum as time reference for the bERPs. Berg and Davies (1988) already stated that the time point zero comparable to ERP research is when the eyelid uncovers the pupils. This happens about 100 ms after the blink maximum and thus qualifies the occipital P200 and N250 as candidates for the P1/N1 complex. The attenuation of the amplitudes compared to the P1/N1 can be explained by superposition of several active brain areas. Based on the interpretation of visual evoked activity in traditional ERP studies, the P200 (P1 in ERP studies) would reflect an exogenous component related to sensory processing of attended incoming visual information which is influenced by various stimulus parameters. The N250 (N1 in ERP studies) then would be related to the allocation of attention to task-relevant stimuli and discrimination of stimulus features (Luck et al., 1990; Luck, 2005).

In the present study, the experimental manipulation did not impact the amplitude of these early components. In contrast, the data of Wascher et al. (2014) revealed an early influence of cognitive effort on the P1 component in the bERP that became more pronounced in the subsequent N2. According to the topography and latency of the N2 as described in Wascher et al. (2014), the comparable component in the present study would be the N250. The absence of a navigation instruction-dependent modulation of these components could then be assumed to reflect comparable mental effort during navigation irrespective of the kind of navigation instruction given.

Concluding, the bERP revealed early components of visual information processing that demonstrated no modulation by the navigation instructions or navigation phases.

#### P300

Berg and Davies (1988) described the posterior P300 of the bERP to be more pronounced when subjects blinked under light conditions as compared to blinking in darkness. In the latter case, the bERP P300 was nearly absent implying that this P300 reflects processing of incoming visual information. The waveform of the component reminds of the well-known P300 at posterior leads in traditional ERP studies and seems to be composed of several activities underlying different cognitive processes. The P300 at posterior leads did not vary systematically between the navigation instruction conditions meaning in turn that underlying higher cognitive processes located at posterior brain areas were comparable for both groups.

#### Late evoked components

Smearing of the ERP at later stages indicates increasing variation of the underlying cognitive and potentially motor processes in time. Thus, the observed sERPs and bERPS revealed no clear components after 600 ms and a slowly returning to baseline activity. This however, is based only on visual inspection of the data and needs further exploration based on more data and using a similar data processing pipeline. This also applies to the late negative level LNL. We are not aware of literature discussing components in eye-movement-related potentials that late and want to point out that especially late phenomena are only comparable to other bERPs if unfolded and preprocessed as described for the present study. If the ERPs are not deconvolved, there is an overlap with subsequent or other physiological events and their related potentials distorting them as explained in Dimigen et al. (2011).

#### Fronto-Central Component

At FCz the bERP was sensitive to differences in the navigation instructions during the navigation phase as reflected in differences from 336 ms to 772 ms comprising the N300 and the LNL. Compared to the baseline phase, the bERP in the long navigation instruction condition was increased about 1 μV while the standard navigation instruction condition remained at the level of the respective baseline bERP. This pattern was visible at several fronto-central leads with increased potentials for the long navigation instructions. At F2 differences in amplitudes became statistically significant during the N300 and at FC1 during the LNL. As this navigation instruction dependent increase was stable and present in most anterior electrodes it is likely due to fronto-central activity reflecting higher cognitive processes evaluating the visual information. In the plots of the raw bERP of Wascher et al. (2014) a peak at 400 ms was visible at Fz and Cz only for the cognitive task but not in a rest condition or the physical task. While the authors did not further discuss this pattern, the present data supports the assumption of higher cognitive processes underlying the amplitude modulation of the bERP around 300 to 400 ms. In fixation-related potential studies investigating visual search in complex real world scenes, a late fronto-central component was revealed in the fERP starting at 300 ms for fixations at a target compared to a distractor face (Kaunitz et al., 2014; Kamienkowski et al., 2018). This component was pronounced at anterior leads while at occipital leads no such difference was visible.

The scalp distribution for the P400 of Wascher et al. (2014) for the cognitive task reveals a similar topography as the fronto-central bERP component in the landmark-based navigation instruction condition during navigation instructions. It seems thus likely that the landmark-based navigation instructions initiated a visual search for the highlighted landmark. Possibly, this visual search and/or information processing of the additional information was more similar to the cognitive task in the Wascher et al. (2014) study while following standard navigation instructions was rather comparable to the physical task of the Wascher et al. (2014) study. This component was less pronounced in the bERP of the walk phase but remained significant in the LNL at FC1. The significant difference at FC3 seems more likely to be due to the attenuation of a baseline difference between the navigation instruction conditions. The significant differences in the fronto-central component were observed dependent on the navigation instruction condition. Increased amplitudes for landmark-based instructions during the instruction phase remained partly visible also during the walk phase. This general tendency points at a cognitive process (e.g. visual search) that is triggered by landmark-based navigation instructions and takes place especially when instructions are given but remains to some extent active during the entire navigation task, even when walking straight segments of the route. This is align with the results of the late positive component shown by Wunderlich and Gramann (2018) in the ERPs of the cued-recall task.

#### Left Parietal Component

In the bERP of the walk phase, a left-lateralized difference in the N250 and P300 components was observed over parietal leads. This modulation was sensitive to the navigation instruction conditions. The long navigation instruction condition showed lower values in the bERP of the walk phase compared to baseline while the standard condition remained at a similar level. The bERP at parietal sites in the study of Wascher et al. (2014) revealed the highest values for rest, followed by the amplitudes in the physical task and most pronounced N2 amplitudes as well as most attenuated P3 amplitudes for the cognitive task. The authors did not discuss lateralized activity but the general parietal focus of the effect is in line with the present results. In Berg and Davies (1988), a slight attenuation of the peaks from a first to a second measurement was noticeable at posterior leads. The second measurement was following directly or 6 min after the first measurement. This would be aligned with a difference between baseline at the beginning of the navigation task and walk phase which comprises most of the blinks in the remaining 40 min. Another explanation for peak attenuation can be the larger total number of blink events in the walk phase in comparison to lower number of baseline blink events. But this does not explain the differences in amplitudes based on the navigation instructions. Concluding, this parietal difference might be due to increasing time on task or landmark-related information processing based on the navigation instructions.

### 4.3 Experimental setup

Table 2 and ERPs especially at occipital leads revealed differences between the navigation instruction conditions already during the baseline before any navigation instructions. Checking all demographic, subjective and individual data collected alongside with the navigation task revealed, that there were some group differences regarding the use of navigation aids despite the random allocation of participants to the groups. P-values of t-tests comparing the group means of the individual measures were below .20 for the 7-point-likert responses to the statements “I use a navigation aid, because I can’t find my way otherwise.” and “If I walk through an unfamiliar city, I know the direction of the start and goal.” as well as in the performance of the Perspective Taking/Spatial Orientation Test (PTSOT, Hegarty and Waller (2004)). This contradicts the assumption of equal means for the different groups.

There might a relation between group differences in individual spatial abilities and the baseline differences of the ERPs at occipital electrode sites. During the baseline, no differences were observed for the first positive component but they emerged with the N200 in the sERP and the N250 in the bERP. These early negative components, if considered to be comparable to the established N1 in visual event-related potentials could reflect different cognitive processes in visual processing of surrounding information affected by individual spatial abilities. Despite baseline differences, an impact of the navigation instruction conditions was shown for assisted navigation after controlling for the baseline potentials.

The absence of a proper baseline is a critical aspect of the present study and the time before the first navigation instruction was chosen as a baseline in a post-hoc fashion. This was the only suitable time window, because this part of the navigation task was expected to be not free of any influence from navigation instructions. The baseline phase started at once with the EEG data collection during the preparation and equipping of the participant and the duration varied between participants. Future experiments should include a controlled and comparable baseline data collection, because especially in between-subject studies, as in the present experiment, group differences in the ERPs independent of the actual experimental the manipulation might be observed. However, by choosing this baseline phase it was still possible to disentangle group differences based on individual measures from those based on the navigation instruction conditions.

### 4.4 Data processing pipeline

The reported data processing pipeline enabled us to successfully extract measures from eye-movement related potentials that are sensitive to changes in visual information processing during free movement in an uncontrolled, real environment. For doing so the only physiological data collected was mobile EEG with 64 channels and one EOG electrode. Blink-, saccade-, and gait-related activity could be extracted applying peak detection in time courses of respective independent components. Overlapping ERPs were deconvolved and controlled for baseline differences using the unfold toolbox.

An important step was the source-based cleaning by separation of brain from non-brain activity. It is questionable whether the IClabel classification is the best way to do this when investigating mobile EEG as the classifier is trained on stationary, laboratory EEG data. The inclusion of new classification criteria, evaluation, and training based on data of moving participants would be necessary to refine the classification quality for mobile EEG. Otherwise IClabel offers a promising approach for automatic source-based data cleaning.

We used the default instead of the lite classifier provided by IClabel because it kept a higher number of brain ICs. On average, two additional ICs reached a classification of at least 30% probability to represent a brain source compared to using the lite version of the classifier. We decided to include as many sources as possible for the sensor-based analysis to include all potential brain contributions. This way the relative importance of each IC for the back-projected data was lower but still with a high likelihood of being caused by brain activity. In a subjective comparison, the lite classifier seemed to be more conservative and more successful in detecting muscle sources and thus should be favored when aiming for extraction of source-based measures (see also the companion paper by Klug and Gramann submitted to this special issue).

Processing the collected mobile EEG data, we realised a strong impact of body movement- and gait-related EEG activity. This impact to the time course of EEG data and the AMICA solution varied between participants. However, especially the 2 Hz shape elicited by the gait cycle clearly compromised the ERPs. Future experiments recording EEG during natural overground or treadmill walking should consider to control for the impact of gait on the recorded signal by adding some kind of motion sensing (see e.g., Kline et al. (2015)).

During the automatic event extraction from the time courses of the respective independent components, percentiles were used as values for thresholds that adapt to the individual signal to noise ratio. When applying the event extraction to other tasks those values might have to adapted.

We focused in our processing and results only on ERP measures. Other research has shown that spectral measures related to blinks can add valuable insights (Wascher et al., 2014; Wascher et al., 2016).

### 4.5 I Conclusion and outlook

The present study demonstrated that EEG activity can be recorded and analyzed in a meaningful fashion using the real world as laboratory. Using blind source separation approaches and subsequent deconvolution of sensor data allowed for extracting brain and non-brain activity that could then be further processed to investigate the impact of navigation instructions on spatial knowledge acquisition. Importantly, using this approach, events representing eye movements like blinks and saccades can be extracted. Those events provide a sufficient number of epochs for event-related potential analyses of the recorded EEG data. This enables new approaches to investigate natural cognition in the real world.

The results in the blink-related results of the present study confirm that the use of landmark-based navigation instructions leads to variations in the accompanying brain dynamics. Differences between auditory navigation instructions are reflected in visual information processing already during the navigation. This difference was mirrored by improved landmark recognition performance of participants that navigated based on landmark instructions.

Future studies will have to replicate the present approach using eye-tracking and scene cameras in combination with high density EEG to establish the link between blinks, saccades and event-related brain activity. More importantly, eye tracking in combination with the scene camera will further provide a valuable source to extract additional events from participants gaze behavior. This will allow for further investigation of the ERP components, comparing eye-movement related ERPs with environment-related events, like fixation on landmark versus fixation at street versus fixation at other traffic participants like pedestrians or cars etc.

In conclusion, the present study demonstrates that it is possible to investigate the human brain in the uncontrolled real world with minimal hardware requirements opening up new avenues for a better understanding of the neural foundation of natural cognition.

## Abbreviations

bERP: blink event-related potential
EEG: electroencephalography
EOG: electrooculography
ERP: event-related potential
fERP: fixation event-related potential
GPS: Global Positioning System
IC: independent component
ICA: independent component analysis
LNL: late negative level
sERP: saccade event-related potential.

## ACKNOWLEDGEMENTS

We would like to thank Prof. Matthias Rötting at TU Berlin for providing the car to transport participants, experimenters and equipment towards and back from the route. Additional we give thanks to Maike Fischer, Christopher Hahn as well as Yiru Chen and the students in the neuroergonomics project course for helping to conduct the experiment.

## AUTHOR CONTRIBUTIONS

K.G. and A.W. designed the research; A.W. collected the original data; A.W. performed the data analysis and drafted the paper; K.G. and A.W. edited the paper.

## DATA ACCESSIBILITY

Data relating to these experiments are available upon request from the corresponding author.

## Footnotes

1 Parameters used for the EEGLAB function pop_rejcont: cleaning based on all electrodes; epoch length of 0.5 s; epoch overlap of 0.25 s; frequency limits to consider for thresholding [10 50]; frequency upper threshold 10dB; four contiguous epochs necessary to label a region as artifactual; once a region of contiguous epochs has been labeled as artifact, additional trailing neighboring regions of 0.25s on each side were added; selected regions were removed from the data; spectrum was computed within the function; hamming was used as taper before fast fourier transformation (FFT)

